# Driving singing behaviour in songbirds using multi-modal, multi-agent virtual reality

**DOI:** 10.1101/2021.10.20.465086

**Authors:** Leon Bonde Larsen, Iris Adam, Gordon J. Berman, John Hallam, Coen P.H. Elemans

## Abstract

Interactive biorobotics provides unique experimental potential to study the mechanisms underlying social communication but is limited by our ability to build expressive robots that exhibit the complex behaviours of birds and small mammals. An alternative to physical robots is to use virtual reality (VR). Here, we designed and built a modular, audio-visual virtual reality environment that allows online, multi-modal, multi-agent interaction for social communication. We tested this system in songbirds, which provide an exceptionally powerful and tractable model system to study social communication. We show that zebra finches (*Taeniopygia guttata*) communicating through the VR environment exhibit normal call timing behaviour, males sing female directed song and both males and females display high-intensity courtship behaviours to their mates. These results suggest that the VR system provides a sufficiently natural environment to elicit normal social communication behaviour. Furthermore, we developed a fully unsupervised online song motif detector and used it to manipulate the virtual social environment of male zebra finches based on the number of motifs sung. Our VR setup represents a first step in taking automatic behaviour annotation into the online domain and allows for animal-computer interaction using higher level behaviours such as song. Our unsupervised acoustic analysis eliminates the need for annotated training data thus reducing labour investment and experimenter bias.

## Introduction

Social communication involves multiple individuals that interact in networks, typically through multi-modal signals, such as vision and sound. Deciphering the mechanisms underlying social communication requires experimental manipulation of the complex multi-modal interactions within the social network. The field of interactive biorobotics provides unique experimental possibilities by letting animals interact with robots to understand, for example, mating behaviours (***Patricelli et al., 2006***; ***Reaney et al., 2008***; ***Partan et al., 2011***; ***Klein et al., 2012***), the underlying rules of shoaling behaviour (***Marras and Porfiri, 2012***; ***Polverino et al., 2013***; ***Kopman et al., 2013***; ***Bonnet et al., 2016***) and communication signals (***Partan et al., 2010***; ***Benichov et al., 2016***). This approach is limited by our ability to build expressive robots that exhibit complex behaviours. What passes for an expressive robot is species and hypothesis dependent, but many animals will readily accept a robot as partoftheirsocial network (***Michelsen et al., 1992***; ***Halloy et al., 2007***; ***de Margerie et al., 2011***; ***Romano et al., 2017***). Building and controlling a small expressive robot might be possible in some cases (***Simon et al., 2019***) but is often not a viable solution for small model animals due to the mechanical and computational complexity involved in fully mimicking natural behaviours.

An alternative to physical robots is to use virtual reality (VR) (***Dombeck and Reiser, 2012***), defined as “a real or simulated environment in which a perceiver experiences telepresence” (***Steuer, 1992***). Current VR setups used in larval zebra fish (***Ahrens et al., 2012***), fruitflies (***Reiser and Dickinson, 2008***) and mice (***Harvey et al., 2009***) virtualise the position of the agent in the environment by providing computer-generated visual feedback. The visual stimulus is generated by measuring the real-world movements of the agent and apply the same translation to its virtual position (***Reiser and Dickinson, 2008***; ***Harvey et al., 2009***; ***Ahrens et al., 2012***; ***Kaupert et al., 2017***; ***Stowers et al., 2017***; ***Cong et al., 2017***). To provide a sufficiently natural virtual environment to interact with and drive the behaviour of an agent, the system needs to be fast enough to analyse, compute and generate the virtual environment within the perceptual real-time of the agent. We refer to this requirement as online operation (***Larsen et al., 2021***). Studying social communication in a virtual environment in most cases also requires multi-modal signals, such as vision and sound and interaction between multiple agents (***Rychen et al., 2021***), but so far VR environments have, to the best of our knowledge, only been used to study single agents. Taken together, to experimentally manipulate social communication, we need a multi-agent VR setup that supports online manipulation of multi-modal signals.

A potentially excellent system for studying social behaviour in a VR context is vocal interaction in songbirds. Zebra finches (*Taeniopygia guttata*) live in societies and form interactive networks through calls (***Zann, 1996***; ***Ter Maat et al., 2014***; ***Anisimov et al., 2014***; ***Benichov et al., 2016***; ***Elie and Theunissen, 2016***). The male song is a learned complex behaviour and is part of the mating ritual where both visual and auditory cues play crucial roles in the natural behaviour (***Zann, 1996***). To situate a zebra finch in virtual reality requires at least sound and vision but is likely also influenced by gaze (***Davidson and Clayton, 2016***) and orientation relative to other agents (***Ljubičić et al., 2016***). Previous work has shown that zebra finches interact vocally with an immobile physical decoy providing audio from a built-in speaker (***Benichov et al., 2016***; ***Benichov and Vallentin, 2020***) and are physically attracted to more life-like actuated zebra finch robots (***Simon et al., 2019***). Furthermore, adult finches can recognize and discriminate between conspecifics from live video feeds (***Galoch and Bischof, 2006, 2007***) and sing song to still images (***Adret, 1997***) or live video feeds of females (***Ikebuchi and Okanoya, 1999***; ***Adret, 1997***). Also, juvenile males can learn song from video and audio playback of a tutor (***Chen et al., 2016***; ***Carouso-Peck and Goldstein, 2019***). Finally, online perturbation of virtual auditory environments can drive active error correction of song (***Sober and Brainard, 2009***; ***Hoffmann et al., 2012***). Taken together, these studies suggest that zebra finches allow studying social behaviour in a VR context. However, no multi-agent VR setup currently exists that supports online, multi-modal manipulation of social communication in zebra finches, and we do not know if zebra finches exhibit normal vocal behaviour when placed in VR environments.

## Results

We present a modular, audio-visual VR environment able to experimentally manipulate social communication. Our system allows for online, multi-modal, multi-agent interaction and focuses on songbirds. Our VR setup is implemented in a box placed inside a cage and the cage is placed in a sound attenuating isolator box. We record and present a high-speed (60 fps) visual environment through a teleprompter system that allows direct eye contact and ensures a realistic visual perspective of the video (Fig 1A). The cage has two perches with presence sensors (Fig 1A); one in front of the teleprompter screen (front perch) and one behind an opaque divider that does not allow visual contact with the screen (back perch). Connecting two VR setups provides the visual impression that the other animal is located 20 cm away (Fig 1B). We furthermore record the acoustic environment and present audio from a speaker located behind the teleprompter to provide the cue that sound and video have the same spatial origin. Data from all sensors is streamed on a network, translated online into events using cloud-based event processing (***Larsen et al., 2021***) and is captured for offline processing by a node connected to storage (historian, Fig 1C). This modular and distributed design allowed for scaling of individual parts of the system (e.g., to add sensors or online software analyses) and can be extended to connect multiple VR setups.

**Figure 1.**
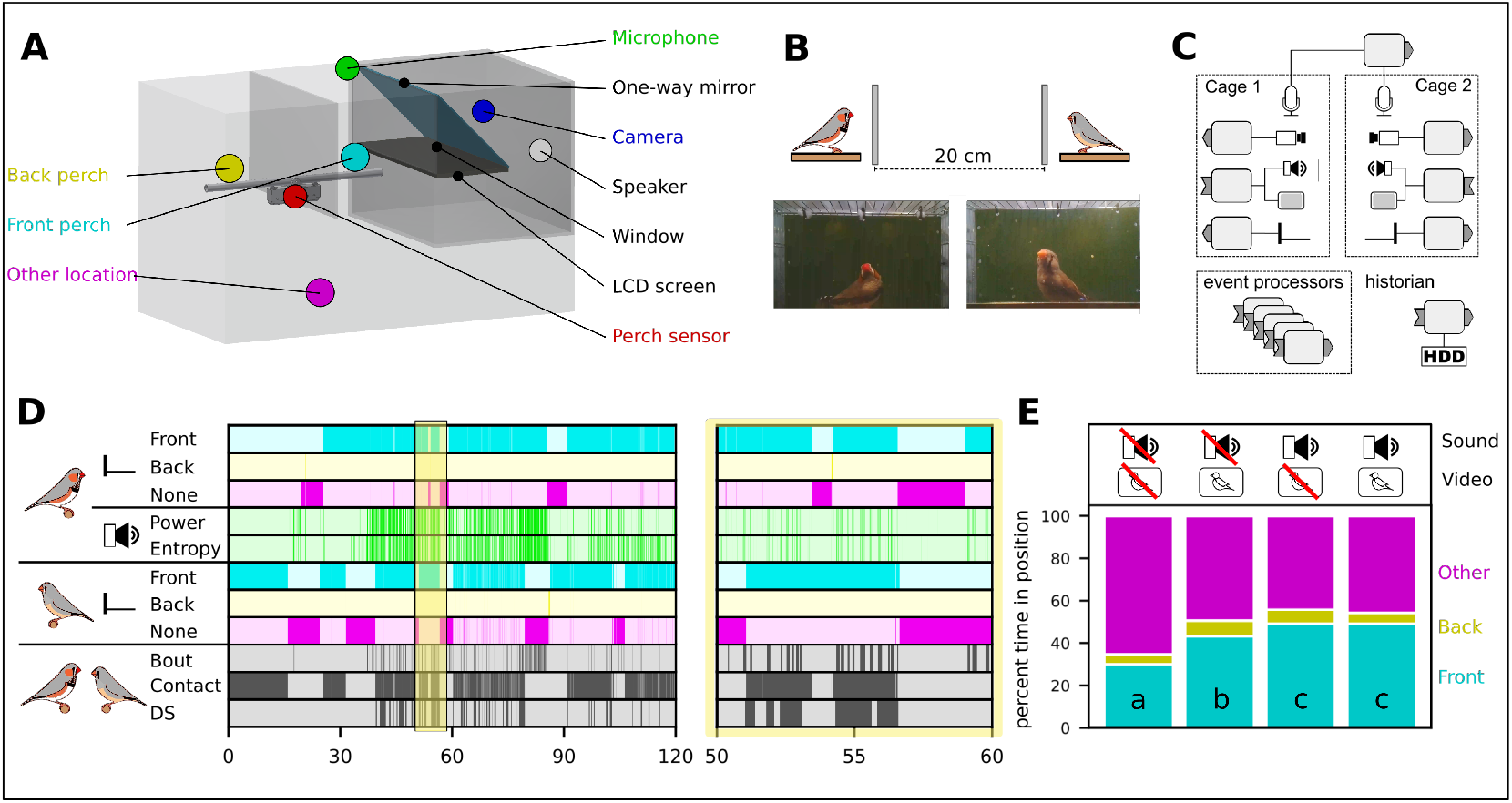
A modular multi-agent multi-modal VR setup for social communication in birds. **A:** The cage equipped with the VR setup and an opaque divider. On the front perch the bird is able to see the screen, while not when it sits on the back perch. **B:** Connecting two VR setups provides the visual impression of the other animal being twice the distance between the bird and the camera away;in this case 20cm. **C:** The distributed hardware architecture of the setup based on ***Larsen et al.*** (***2021***). See Methods for more detail. **D:** Perch and acoustic events produced during two hours of communication through the VR setup and a 10-minute zoom of the area with yellow overlay. For complex event definitions see main text. **E:** The perch preference in four different experimental conditions. Different letters denote significant difference (two-proportion z-test, 1 % significance level, n=24).

Event Processing (***Cugola and Margara, 2012***) was used to represent onset and offset of behavioural features and event streams from multiple producers were combined to form new events (Fig 1D). The position of a single bird generated three different events for absence/presence on the front or back perch or other location (Fig 1D, cyan, yellow and magenta lines). When connecting two VR setups with one bird in each cage, a visual contact event was defined as both birds perched in front of the screen and thus able to see each other. To detect and identify vocal signals, the audio stream from the male was analysed online (see methods) to generate events based on power and entropy threshold-crossings (Fig 1D, green lines). A bout of song was detected by combining power and entropy with hysteresis thus suppressing most noise. Finally, a directed song (DS) event was generated when bout and contact were active at the same time (Fig 1D, bottom line) in other words, when a male was singing while both male and female were sitting on the front perch.

To investigate the animals’ motivation for social interaction through the VR setup, we measured the perch preference of twelve pair-bonded male-female couples in four different audio-visual modality combinations of speaker/screen on and off. Our data showed that birds spent significantly more time on the front perch when one modality (either sound or video) from the other bird was on (Fig 1 E). When both video and audio modalities were on, birds also spent more time on the front perch than video-only but not when compared to live audio (two-proportion z-test, 1 % significance level, n=24). This demonstrates that the birds were attracted to both audio and visual signals of another individual supplied by the virtual reality system.

To demonstrate that the VR setup provided a sufficiently natural environment for social communication, we exploited two key behaviours in communication between pair-bonded individuals: call timing and directed song. Coordinated call production between partners is a well described behaviour in birds, where it is thought to influence pair-bond maintenance and mate guarding (***Elie et al., 2010***). Zebra finches show time-locked call behaviour using two types of calls: Tet and stack calls (***Ter Maat et al., 2014***). Both are short, low power vocalizations used when the birds are physically close together (***Zann, 1996***; ***Elie and Theunissen, 2016***). We quantified the call timing of stack calls between established pair bonded couples communicating through our VR environment (Fig 2A) and identified calls using a supervised random forrest classifier (see Methods). With both audio and visual modalities on, the delay from hearing a call to producing one was unimodally distributed (Fig 2B) with a peak delay at 291 ms (median: 271 ms, range: 231-431 ms, N=8, Fig 2BC). This data is consistent with call timing delay measured between free-moving pairs in colonies (Ĩ91 ms (***Ter Maat et al., 2014***), 249-466 ms (***Benichov et al., 2016***) and 68-283 ms (***Anisimov et al., 2014***)). Because calls were synchronized and contingent on the call of the mate, we conclude that the birds displayed natural call timing behaviour through the VR setup.

**Figure 2.**
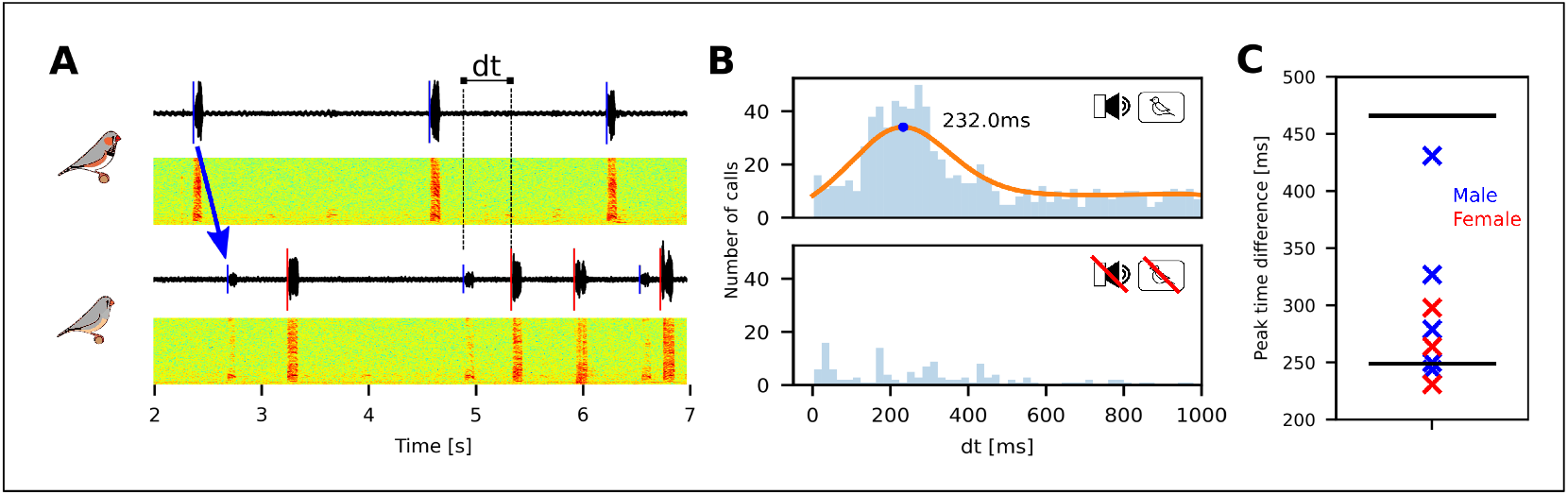
Zebra finches communicating through the VR setup exhibit natural call timing behaviour. **A:** Sound oscillograms and spectrograms for the stack calls of a male and a female communicating through VR setups. Detected onsets are indicated by vertical lines on the oscillograms. The arrow denotes that the playback of the male call can be seen in the recording of the female. We measure the time dt from playback to answer. **B:** Histogram of the elapsed time between the playback of the mate’s call until next call in one pair. The peak of the kernel density estimate is marked. Bottom plot shows a histogram of time differences for the same pair with the system off. **C:** Summary showing the peak for all 8 birds. The black lines show the range of values reported in (***Benichov et al., 2016***).

Next, we studied whether the VR setup provided a sufficiently natural environment for males to exhibit natural singing behaviour to their virtual mate. Male zebra finches sing both to females (directed song, DS) and not directed towards any particular conspecific (undirected song, US) (***Zann, 1996***). The song consists of introductory notes and a stereotyped sequence of syllables, called the motif, that is often repeated several times to form a song bout (***Zann, 1996***; ***Sossinka and Böhner, 1980***). Although the DS and US motif consist of the same syllable sequence, several key acoustic features are different between DS and US. The DS motif is delivered faster and is preceded by more introductory notes ***Jarvis et al., 1998***). It also has more repetitions of the motif in each bout, increased sequence stereotopy (***Sossinka and Böhner, 1980***) and DS syllables exhibit less variation in the fundamental frequency of harmonic stacks (***Kao et al., 2005***).

We studied five established pair-bonded couples communicating through our VR environment and we isolated candidate DS events as the simultaneous occurrence of bout and contact events, i.e., when both animals were perched in front of the screen and the male was vocalizing. The video segments of potential DS events were subsequently scored for accompanied behaviour by experienced observers (IA, CPHE). All (5/5) males sang directed song to their virtual mates and displayed courtship behaviours, such as fluffing, beak wipes and jumping, that are indicative of DS (Fig 3A; Movie M1) at high intensity. The coefficient of variance of the fundamental frequency (Fig 3BC) was significantly lower (Wilcoxon signed rank test, 5 % significance level, n=5, Fig 3D) when the VR system was on compared to off further indicating DS. Taken together our data strongly suggest that all males sang DS to their virtual mates.

**Figure 3.**
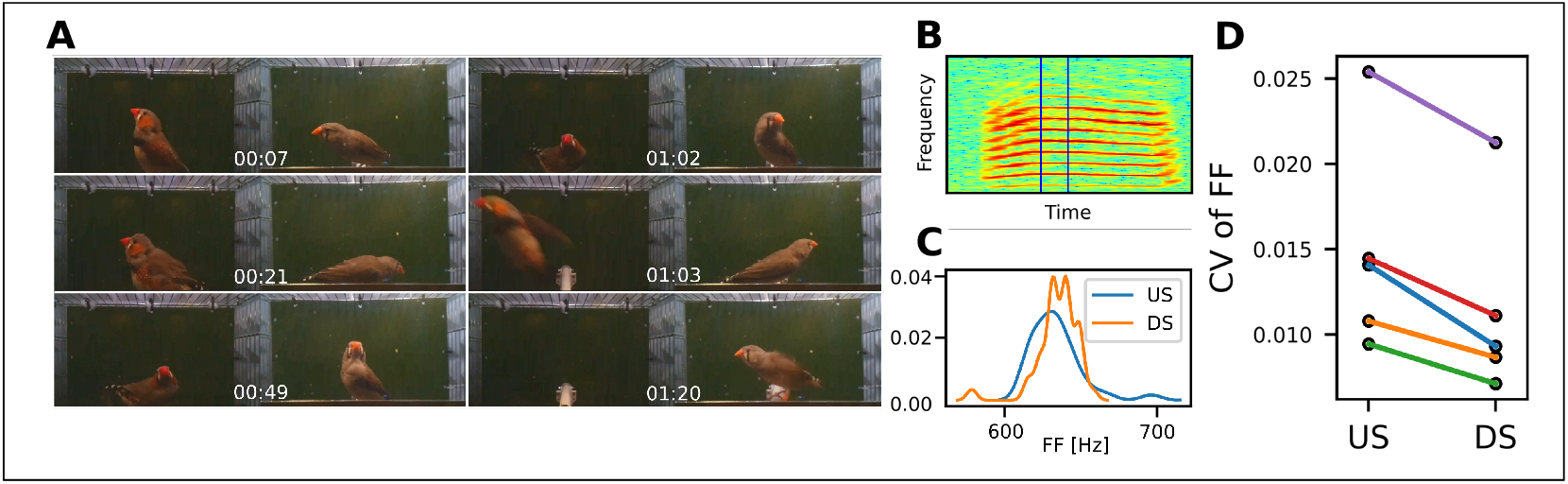
Adult male zebra finches sing directed song to their mate through the VR system. **A:** Male (left) and female (right) displaying behaviours associated with directed song. The full video is available in the supplementary materials (Movie M1). **B:** Spectrogram of a stack call used to estimate fundamental frequency. The vertical lines indicate the part of the syllable used to estimate fundamental frequency **C:** Kernel density estimates for the fundamental frequency of the syllable above based on 30 renditions with the system on (DS) and 30 with the system off (US). **D:** The Coefficient of Variance of the fundamental frequency is significantly lower when the VR system is on (Wilcoxon signed rank test, 5 % significance level, n=5) indicating DS.

In summary, call timing between individuals was comparable to that of freely communicating animals and males sang DS to their virtual mates, which showed that our VR system provided a sufficiently natural environment for multi-modal, multi-agent social communication in songbirds.

A final crucial component in the design of an online VR system is the ability to manipulate an agent’s environment and thereby drive its behaviour within its perceptual real-time. When zebra finch males sing DS to a female, they typically habituate to its presence, which leads to a reduction in the number of motifs per minute (***Jarvis et al., 1998***). In experiments requiring extended periods of DS and/or a high number of motifs, this effect is typically countered by introducing novel females to reinvigorate the male (***Jarvis et al., 1998***; ***So et al., 2019***). Here we aimed to drive DS behaviour and increase the number of produced motifs by presenting different virtual females based on the online measured song performance of the male.

Driving song behaviour based on song performance requires online detection of the stereo-typed syllable sequence, i.e., the motif. Therefore we developed a novel, unsupervised online motif detector (see Methods). The detector is based on dimensionality reduction of feature vectors generated from the spectrogram of sound segments (Fig 4A-E). Training of the model was based on 60,000 feature vectors per animal, each representing a segment of sound. The feature vectors were embedded in a 2D space using t-distributed Stochastic Neighbor Embedding (***Maaten and Hinton, 2008***) and the watershed transform (***Meyer, 1994***) was used to cluster the space into a behaviour map (***Berman et al., 2014***). Next, we computed the transition probability matrix between all syllables in the training data (Fig 4H) and used it to detect the most stereotyped sequence of syllables by starting at the globally most likely transition and following the path of locally most likely transitions (Fig 4H). In all the males, this path contained a cycle that we defined as the motif of the individual and it was confirmed by experienced observers (IA, CPHE) to be the correct motif.

**Figure 4.**
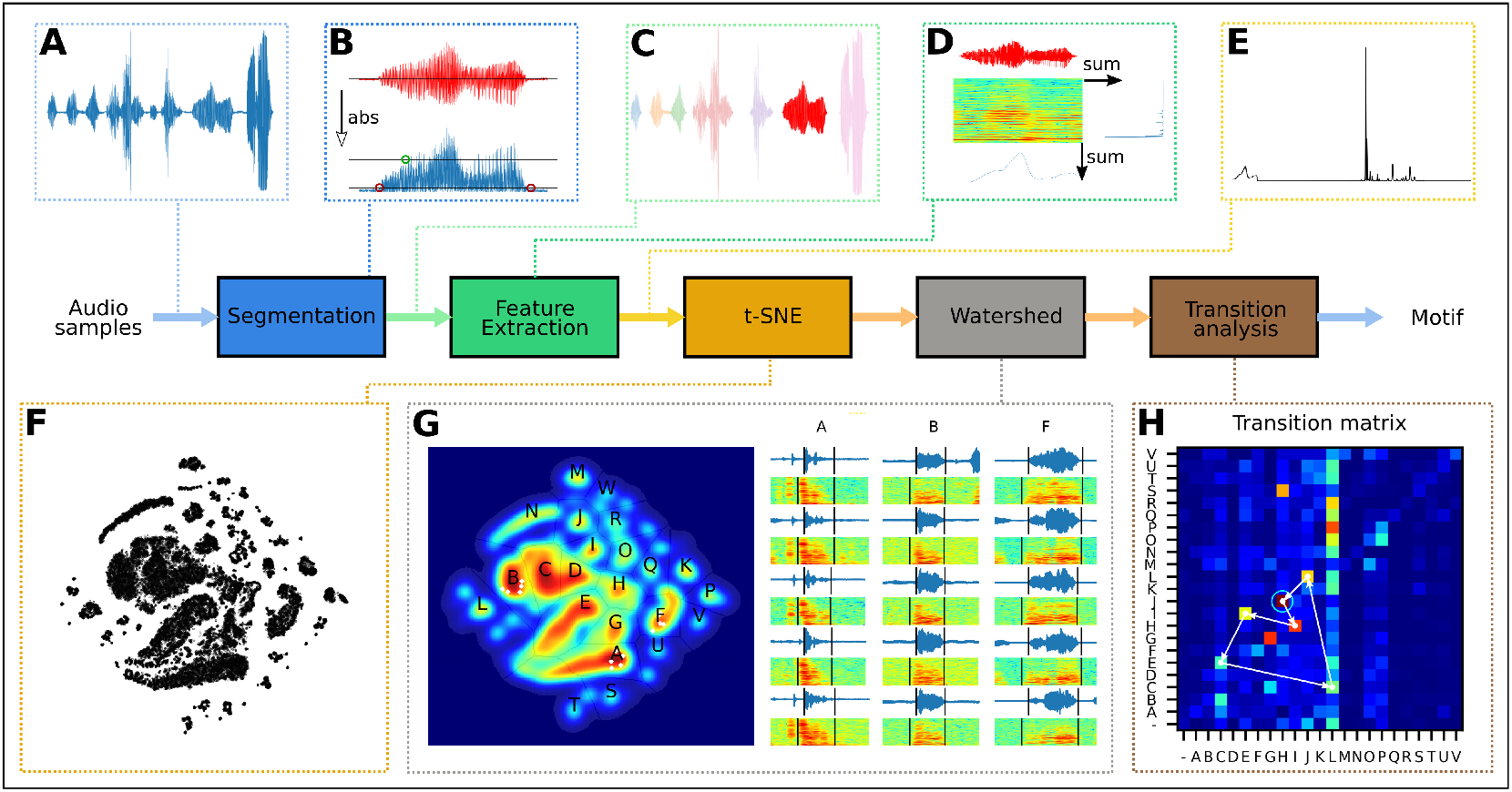
Unsupervised training of motif detector pipeline. **A:** The raw sound signal is received as a continuous stream. **B:** When the absolute value of the samples crosses the high threshold, we search back and forth for onset and offsets based on crossing of the low threshold. **C:** The segmented signal. **D:** A spectrogram of the sound is generated and two vectors are computed by summing the rows and columns, respectively. **E:** The two vectors are normalised and concatenated to a 746-dimensional feature vector. **F:** Dimensionality reduction of 60k feature vectors into a 2D space shows that sound segments cluster together. **G:** Regions of stereotyped sounds are labelled. Examples of randomly picked sounds from three different regions are indicated by white dots on the density plot and with oscillograms and spectrograms on the right. Each column is a different region. **H:** The most common song motif is extracted by forming the transition probability matrix and following the most likely transitions forming a loop. The loop represents the most stereotyped sequence and is defined as the motif.

Next, we extended the method to detect motifs online (Fig 5A). We detected syllable events by analysing the audio stream and post-embedding the sound segments into the previously computed 2D space (Fig 5B). Syllable events were then collected in sequence events that were screened for ordered subsets of the motif (see Methods) to create the motif event (see example in Fig 5C). The entire process was parallelised to achieve online detection.

**Figure 5.**
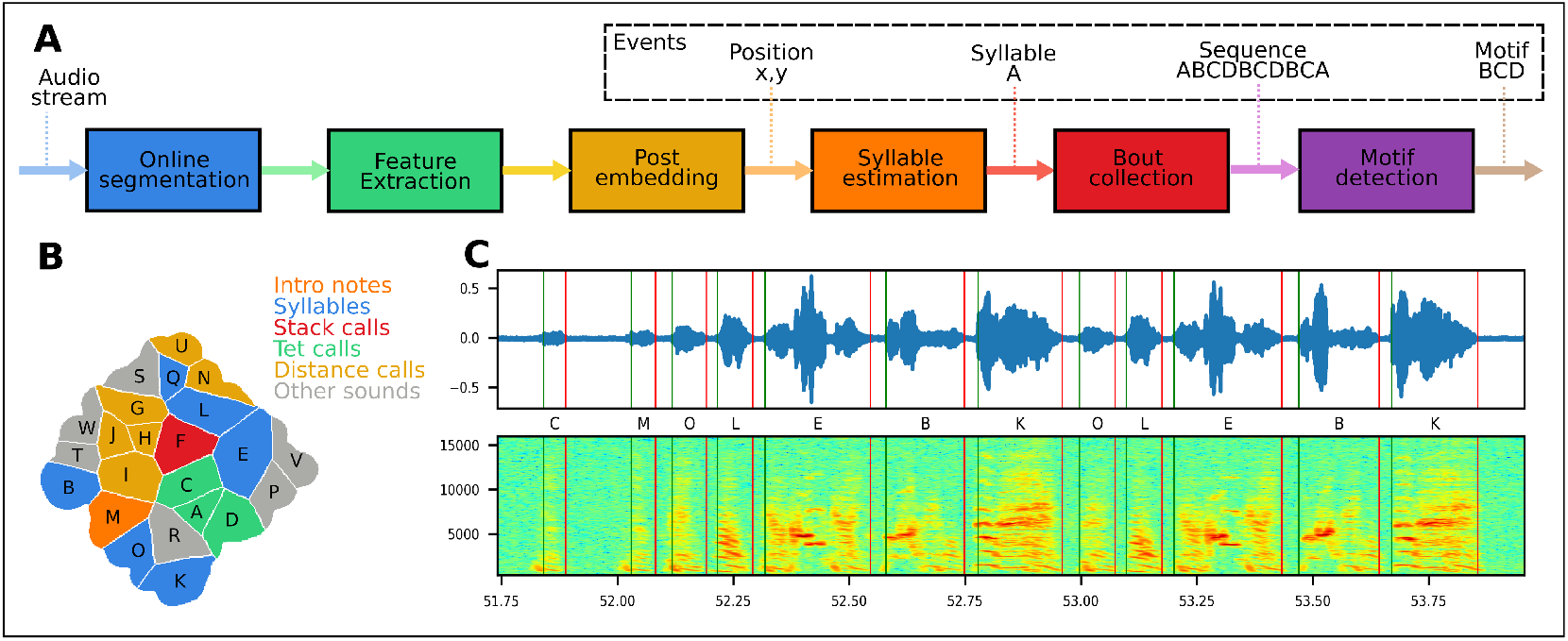
Pipeline for online analysis and event generation. **A:** The pipeline for online analysis uses the methods described in fig. 4A-E for segmentation and feature extraction. The resulting feature vector is post-embedded in the 2D space and classified based on which region it falls within. Syllables are collected in sequences and screened for motifs. **B:** The behaviour map annotated manually based on sample sound segments from each region **C:** Song spectrogram with vertical lines indicating onset (green) and offset (red) events of automatically detected syllables. The letters indicate the class of the syllable.

Next, we exposed males to one minute audio-visual recordings of one female in an excited state from the DS experiments, as long as motifs were detected (see Methods). After three minutes without motif detection, we switched to the audio-visual recordings of another female. Driving the behaviour over two hours, males sang significantly more motifs (Wilcoxon signed rank test, 5 % significance level, n=9) compared to the control period (Fig 6AB). To confirm that the birds sang DS motifs, we computed the CV of FF in a motif syllable containing a harmonic stack. The CV was significantly lower during the driving period compared to the control period (Wilcoxon signed rank test, 5 % significance level, n=6), which strongly suggest that the males sang DS to the virtual females (Fig 6C). Taken together, our VR system made the birds sing more directed motifs in two hours compared to undirected motifs in the control, thus demonstrating the ability to drive directed song behaviour.

**Figure 6.**
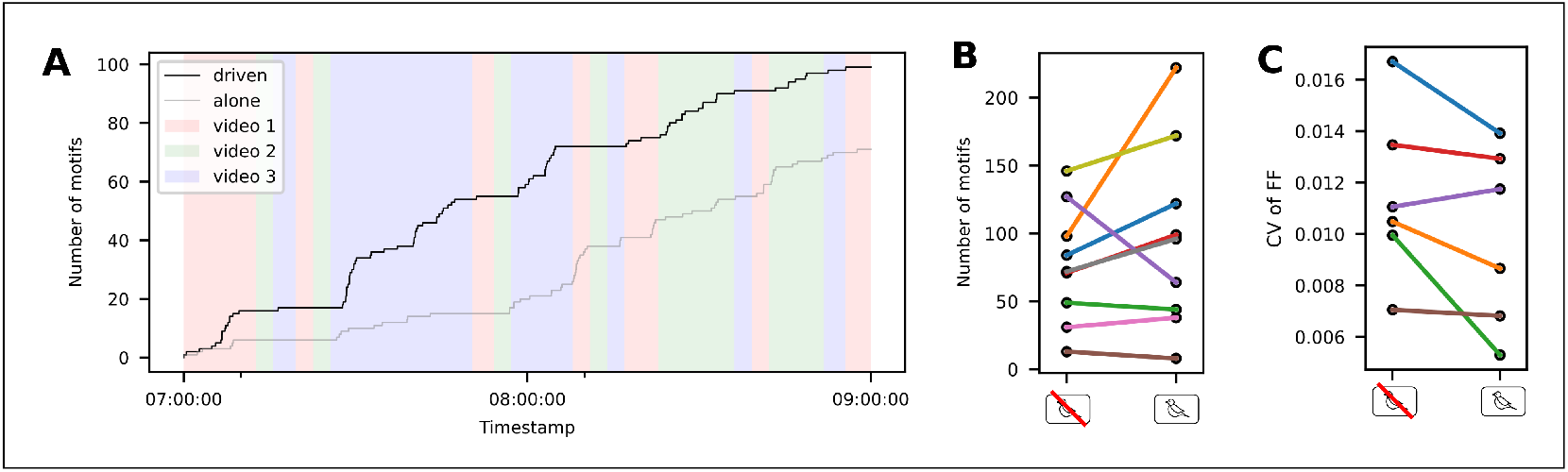
Number of directed motifs can be increased with the help of online motif detection. **A:** Cumulative sum of detected motifs for one bird during a two-hour period of driving the behaviour by switching virtual female individuals and the same two hours on the following day without video. Background colours show which video was playing during that time. Videos change dynamically based on number of motifs. **B:** The number of motifs sung was significantly higher (Wilcoxon signed rank test, 5 % significance level, n=9) during the two hours with video compared to the two-hour control without video demonstrating that the video drives them to sing more. **C:** The coefficient of variance for the fundamental frequency was significantly lower (Wilcoxon signed rank test, 5 % significance level, n=6) during the two hours with video compared to the two-hour control confirming that they sing directed song to the video.

## Discussion

We present a VR environment to study social communication that allows online, multi-modal, multiagent interaction. Zebra finches communicating within the modular VR environment emitted calls that were synchronized and contingent on the call of the mate with response latencies as in real life situations (***Benichov et al., 2016***; ***Ter Maat et al., 2014***; ***Anisimov et al., 2014***). Furthermore, our data show that males exhibited high-intensity courtship behaviour and sang directed song to their virtual females. To detect DS events, we used an easily implemented definition of DS as song that occurs while the birds had visual contact. Previous studies also defined DS as song when the male was singing oriented towards a conspecific female (***Adret, 1997***; ***Chen et al., 2016***), but did not confirm this classification by further acoustic analysis such as decreased DS motif duration (***Jarvis et al., 1998***), or decreased variation in the fundamental frequency of harmonic stacks in DS syllables (***Kao et al., 2005***). ***Ikebuchi and Okanoya*** (***1999***) classified each song rendition as either directed or undirected based on dance behaviour but did not indicate their criteria for this classification. Using both behavioural and acoustic analysis we confirmed that song elicited under our definition was indeed DS. Taken together, these data strongly suggest that the VR environment is sufficiently realistic to elicit the full spectrum of courtship behaviours.

We present and implemented a syllable-based unsupervised audio classifier which we think will be widely applicable in bioacoustics. Unsupervised clustering methods have been used in the analysis of vocalisations (***Tchernichovski et al., 2000***) but are typically based on a few dozen acoustic features. A more data-driven approach is to use spectrograms directly as high dimensional features (***Kollmorgen et al., 2020***), which however imposes extensive computational costs. Here, we compressed the spectrograms to arrive at a manageably sized feature vector thereby keeping computational costs low. Especially for stereotyped behaviours, unsupervised methods like t-SNE (***Maaten and Hinton, 2008***) excel because clusters of repeated behaviours stand out from noise (***Berman et al., 2014***). Furthermore, we parallelized a variation of the post-embedding algorithm described in ***Berman et al.*** (***2014***) to achieve online classification. Lastly, we could determine each individual’s motif in an unsupervised way by assuming only that the motif is the most repeated syllable string, thus exploiting the fact that zebra finch song is highly stereotyped. Our unsupervised method eliminates the need for annotated training data and thereby reduce labour investment and the risk for experimenter bias.

Our setup represents a first step in taking automatic behaviour annotation into the online domain. We used events to represent behaviour and event-processing and microservices to achieve online capabilities (***Larsen et al., 2021***). Several studies have demonstrated the power of online processing in closed loop assays in neuroscience (***Grosenick et al., 2015***; ***Nourizonoz et al., 2020***), to manipulate pitch in songbirds (***Lohr et al., 2003***; ***Brumm and Slabbekoorn, 2005***; ***Riedner and Adam, 2020***) or to provide virtual reality (***Ahrens et al., 2012***; ***Reiser and Dickinson, 2008***; ***Harvey et al., 2009***). Those studies take advantage of computationally attractive features, such as action potentials or acoustic features, to allow near real time system response. Our approach is slower but allows system response to target higher levels of behavioural organisation, such as syllables or motifs in the audio domain. The limiting factor is the ability to infer online the behaviour of the animal, in other words to parallelise and optimise algorithms for behaviour annotation.

Our VR system is modular and can be extended to multiple setups or to add more sensors, actuators, and computational units. We expect our online, modular setup to be applicable to other species of social birds and mammals. We deliberately based the setup on cheap distributed computers, free and open-source software, and cloud computing to ease the reuse of hardware and software modules in other projects, and make it easier for multiple developers to contribute. The distributed architecture complicates the system and increases the minimum latency, but allows it to scale linearly, makes it easier to maintain, and makes it resilient to single node failures. The system can be deployed anywhere with network and thereby enables global-scale social communication experiments. VR setups situated around the globe could thus be connected and allow for unique long-term communication experiments between labs that are physically far apart.

## Methods

### The VR setup

The VR-setup was built on the teleprompter principle, where a slanted one-way mirror allows the camera to record the bird through the mirror while the bird sees the reflection of a screen below (Fig 1A).

A microphone is placed outside the cage above the perch in front of the screen while the rest of the system is placed inside a painted wooden box placed in the cage. The one-way mirror is constructed from a sheet of 3 mm transparent acrylic plexiglass coated by 0.02 mm silver one-way film with 70 % light admittance and 99 % reflectance.

The screens are trichromatic 7” LCD displays in 800×480 pixel resolution. Although birds possess at least tetrachromatic or even pentachromatic vision (***Emmerton, 1983***), previous studies showed that males sing when presented with live video of conspecific females on trichromatic screens (***Adret, 1997***; ***Ikebuchi and Okanoya, 1999***; ***Galoch and Bischof, 2006***). However, critical to eliciting courtship behaviour was the use of 100 Hz screens (***Ikebuchi and Okanoya, 1999***; ***Galoch and Bischof, 2006***) that are above the flicker-frequency of birds (***Emmerton, 1983***; ***Nuboer et al., 1992***) or non-flickering liquid-crystal displays (LCD). Therefore, we decided to use 60 Hz LCD screens that present slower, but continuous, flicker-free images to the birds. The video is recorded with a Raspberry Pi Camera V2 in 800×480 pixel resolution at 60 frames per second (fps) and streamed to the network from a Raspberry Pi 3. The video delay was measured by simultaneously turning on an LED in both boxes and recording a video with an external camera showing both the LED and the screen. By counting the number of frames from the local LED turns on to the remote LED from the other box can be seen on the screen the delay can be calculated. This delay was measured to 383 ms (23 frames at 60 fps).

The audio playback comes from a 1.5 W mini-speaker placed behind the mirror and had to be slightly attenuated (−6dB) to avoid acoustic feedback. The sound was recorded and streamed from a multi-channel recording array (***Andreassen et al., 2014***) using Knowles FG23329-PO7 microphones. The recording equipment is not part of the developed VR setup and it could be replaced by any system capable of streaming audio. The audio delay was measured by making a loud sound (with a clicker) in one box and timing the difference between that signal in one box and the version played back in the other box. This delay was measured to be 308 ms ± 4 ms.

Figure 1 C shows the architecture of the system. Each VR-box contains two Raspberry Pi 3 model B connected to a gigabit switch. One is connected to the camera and is only responsible for streaming video. The other, connected to display and speaker, is responsible for playback of sound and image. A third Raspberry Pi 3 is placed on top of the cage responsible for polling the perch sensor at 20 Hz and emitting state changes as events. It also measures temperature and humidity in the box and emits those as events every minute. The multi-channel microphone array is placed outside the isolator box with a microphone placed in each cage. All computers on the network are synchronised to within milliseconds using the Network Time Protocol (***Martin et al., 2010***) implemented with chrony (***Lichvar, 1999***).

Data is streamed to IPv6 multi-cast groups following the publish-subscribe pattern (***Birman and Joseph, 1987***). A PC in the bird room acts as historian, saving the data streams. Data is offloaded to a Ceph (***Weil et al., 2006***) persistent storage cluster placed in our data centre. Several event processors continuously analyse the data streams, producing events. These are running in a docker swarm (***Merkel, 2014***) cluster also in our data centre.

The two-layered architecture is based on data streams and event streams (***Larsen et al., 2021***). An event is an association between a specific time and a specific property, in this case a behaviour. A data stream contains sampled data from sensors such as cameras and microphones while an event stream consists of events produced by data stream processors or by asynchronous sensors such as contacts.

The continuous audio stream is analysed online to produce the power and entropy events. This analysis is based on estimating the power by squaring the sample values and the entropy as the ratio of the geometric mean to the arithmetic mean. The analysis is implemented as plug-ins to the media-streaming framework gstreamer (***Gstreamer, 2001***). The estimates are thresholded with hysteresis (***Larsen et al., 2021***) and published as kafka events (***Vohra, 2016***). A perch sensor installed in the cage directly generates a perch event every time the bird changes location in the cage. Based on those three events, three complex events are generated, namely bout, contact and directed song. The bout event is active when both power and entropy events are active, and the contact event is active when both birds are perched in front of the screen. The directed song event is active when the bout and contact events are active (Fig 1 D). Event processing was implemented as microservices in docker containers (***Merkel, 2014***) for high modularity and was running in our data centre.

### Animals and husbandry

Adult male and female zebra finches (*Taeniopygia guttata*) were kept pairwise in breeding cages at the University of Southern Denmark, Odense, Denmark on a 12 h light:dark photoperiod and given water and food ad libitum. All experiments were conducted in accordance with the Danish law concerning animal experiments and protocols were approved by the Danish Animal Experiments Inspectorate (Copenhagen, Denmark).

We used adult zebra finches (> 100 days post hatch) that were established breeding pairs (meaning that they had produced at least one clutch of offspring together) in the animal-animal communication experiments and additionally also single males for the animal-computer experiments. When not in experiment, the birds were kept pairwise in breeding cages or in aviaries containing hundreds of individuals. Under experiment the birds were isolated in sound-attenuated boxes for a maximum of ten days before returning to their usual surroundings. The birds had access to food and fresh water ad libitum served at the bottom of the cage and from feeders at the side of the cage. In the VR setup, the birds were kept on a 12 h light:dark photoperiod. The temperature was kept between 22 and 28 °C and the relative humidity at 50-60 %. The temperature difference between the position in front of the screen (front perch) and behind the blind (back perch) was measured with the system fully on to be 0.4 °C (± 0.3 % accuracy). The isolator boxes attenuated sounds in the 200-8,000 Hz range by 40dB (measured by playing back sound in the isolator and record sound levels both inside and outside the box. A fan ensured air flow in the box and provided cooling for the equipment located inside.

### Sound segmentation

Segmentation was based on the silence between syllables and was calculated from the amplitude of the signal (Fig 4B). We used two threshold values for discriminating between sound and silence. The input signal (Fig 4A) was normalised to range but otherwise not pre-processed. Starting from the *i_on_*th sample where the absolute value of the sample *s_i_* surpasses the on-threshold *t_m_* (0.5)

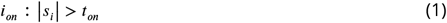

we searched backwards in time to find the onset sample number *i_onset_* defined as the sample where the peak-to-peak amplitude over *w* samples (325) was below the off-threshold *t_off_* (0.5).

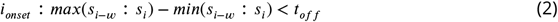

Similarly we searched for the offset sample number *i_offset_* as new samples arrived.

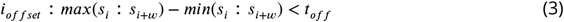

As soon as the last sample was received the segment was passed on to the next stage of the pipeline. Segments shorter than 30 ms or longer than 300 ms were discarded since the duration of zebra finch syllables is expected to be within that range. We implemented both an online and offline version of this segmentation algorithm and used it for all the experiments presented in this paper.

### Perch preference protocol

The same data was used for all the animal-animal communication experiments. The VR-setup was powered down for at least 2 hours before the birds were moved to the isolator box (day 0) and left off for at least 24 hours before it was turned on for another full day (day 1). Experiments ran on the following days starting when the cage lights were turned on and for two hours thereafter. Day 2 was always with black screen and no sound and the following days the system cycled through perturbations of two speaker states (off, on) and two screen states (off, on) in randomised order. After the experiments the birds were returned to their home cages. To investigate the motivation for using the VR setup, we looked at the perch preference in different states of the system. Based on the perch sensor, an event was emitted every time the bird changed position in the cage and summing the duration of the events gives a measure of the proportion of time spent in each position (Fig 1E).

### Call-timing protocol

The audio was segmented as described above and combined into one big dataset covering 12 hours a day for all 12 pairs. To provide training data, we then hand-annotated for each bird the first 30 minutes with both video and audio on. To ease annotation, we used pre-clustering based on cross-correlation, so the observer was presented with oscillograms, spectrograms and sound from one minute at a time that had already been clustered into groups of sounds with high crosscorrelation maximum. The observer then had to name the groups and correct mistakes made by the pre-clustering algorithm. The classes found were song, stack calls, distance calls, echo (loud sounds from the other bird triggering the segmentation), wing flapping and noise. Classification of vocalisations followed the descriptions in ***Zann*** (***1996***). Based on the annotations a random forest classifier (***Breiman, 2001***) with 100 estimators was trained for each bird ranging in mean accuracy (10 % hold out) from 0.82 to 0.96. To investigate call timing, we measured the time difference from the playback of a stack call (onset + delay) to the next stack call emitted by the animal of interest up to a maximum of 2 seconds. Histogram of the time differences were constructed (200 bins) and plotted with Gaussian Kernel Density Estimates (KDE, bandwidth=100, fig 2B).

### Directed song protocol

To confirm directed song in the animal-animal communication experiments, we selected videos with potential female directed song based on the definition that both birds were on the front perch and the male was vocalising. The videos were then scored by experienced observers (IA, CPHE) for the display of hopping, jumping, beak wiping, looking at the mate and fluffing plumage (see figure 3D for examples). As a quantitative measure, we calculated the coefficient of variance of the fundamental frequency, which is lower in DS compared to US (***Kao et al., 2005***). However, this measure is extracted from stack syllables without frequency modulation such as the one shown in figure 3B. The motif of five birds contained a suitable syllable. From spectrograms of the motifs identified by the online motif detector, we manually selected the same place in the stack syllable (Fig 3B) in 30 motifs from each individual. The fundamental frequency was estimated from 2048 (42 ms) samples using the YIN algorithm (***De Cheveigné and Kawahara, 2002***).

### Animal-computer communication experiments

For the driving experiments, the male was left to habituate to the new surroundings until he produced at least 10 motif repetitions during the first two hours after lights on (day 0). On the following day (day 1) we ran the driving experiment, meaning that videos were displayed showing excited females. Three videos of different females were used, each one minute long, taken from the animal-animal communication experiments. In case of pair-bonded males, the established mates of the focal animals were not among the female videos. In case of single males the 60,000 training samples were recorded over three days prior to the experiment. The logic of the system is that every time a motif is detected, a timer is reset. If the timer ran out (3 minutes since last motif) the next video was displayed and otherwise the same video kept getting looped. On day 2 we recorded the control without video playbacks.

### Online motif detector

The feature extraction is based on summing the rows and columns of the spectrogram (Fig 4D) and concatenating them to form a feature vector (Fig 4E).

First a spectrogram of the segment is formed by applying Short-Time Fourier Transform (STFT) with FFT size of 1440 and stride of 25 samples. The parameters are all based on the sampling frequency of 48 kHz, the duration of sounds (30-300 ms) and the desired number of time bins. The smallest spectrogram we can make has just one time bin and thus the maximum FFT size is:

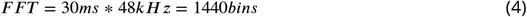

The stride parameter can then be calculated:

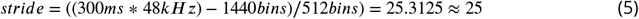

The spectrogram is cropped to the approximate audible range for zebra finches 200 Hz to 8 kHz (234 bins) and the time dimension is cropped to the first 512 time bins corresponding to 300 ms (zero-padded if the segment duration is shorter). The rows and columns of the spectrogram are summed and the two resulting vectors *F_t_* and *F_f_* are concatenated to form a 746-dimensional feature vector *F* (Fig 4E).

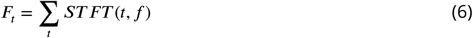

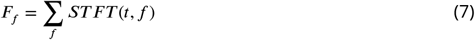

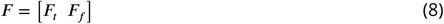

Each vector is normalised to have a sum of one before concatenation.

A training set is created consisting of 60,000 feature vectors from the same individual, each representing one sound segment. We embed each of these high-dimensional points in a two-dimensional space (Fig 4F) using the t-SNE method introduced in ***Maaten and Hinton*** (***2008***). The method minimises the relative entropy between two distributions, one representing the high-dimensional points and one representing the low-dimensional points, so that close points in the high-dimensional space are also close in the low dimensional space.

Since we are interested in stereotyped behaviour, we then loosely follow the method described in Berman et al. (2014) placing a Gaussian kernel (bandwidth=15) on each embedded point we generate a density plot (Fig 4G), and we find all peaks that are separated by a distance of 15 or more. Using the watershed algorithm (***Meyer, 1994***) on the inverted density plot, we get a set of clusters, each representing roughly a stereotyped syllable. Examples from three different regions can be seen in figure 4G. The further a point is from the peak of the region the more likely it is to be mis-classified and thus distance from peak could be used to indicate certainty of the classification. We found that some regions represent a merge of two syllables while some represent part of a split syllable. For higher accuracy in detecting the syllables this information could be used for post processing or better means of segmentation could be introduced.

After the training phase a new data point *z* is embedded based on the already embedded points, largely using the method described in ***Berman et al.*** (***2014***) appendix D. The perplexity parameter of the t-SNE algorithm can be interpreted as a measure of the number of nearest neighbours (***Maaten and Hinton, 2008***) and therefore we only consider the ‘perplexity’ nearest points *X* in the high dimensional space found using the ball tree algorithm (***Omohundro, 1989***).

We then choose an embedding *Z* of the new point *z* such that conditional probabilities in the low-dimensional space *q_j|z′_* are similar to those in the high-dimensional space *p_j|z_*. The conditional probability of a point *x_j_* ∈ *X* given the new point *z* is:

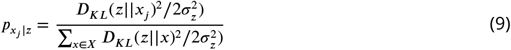

where *X* is the vector of nearest points in the high-dimensional space, *z* is the new point in the high-dimensional space, sigma is found by a binary search for the value that produces a conditional probability with the perplexity set by the user and *D_KL_* is the relative entropy given by:

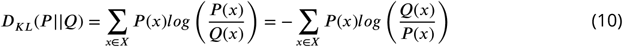

The conditional probability of a point 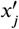 in the low-dimensional embedding *X′* given the new embedding *z*′:

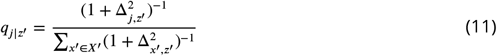

where Δ_*a,b*_ is the euclidean distance between the points *a* and *b*. Since *z′* is the only unknown, we can find it by minimising *–D_KL_* between the conditional probability distributions:

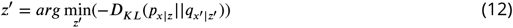

using the Nelder-Mead simplex algorithm (***Nelder and Mead, 1965***) and a start guess being the centroid of the embedding *X′* of the nearest points *X*. If the start guess is not in the basin of attraction of the global minimum, it means that the new point is not like any points presented during training and the embedded point will shoot towards infinity (***Berman et al., 2014***).

One instance of the segmentation algorithm was running for each of the two audio channels used in the experiment and a new feature vector was formed for each detected segment and placed in a queue. A pool of 6 workers (containers running in the cluster) processed feature vectors from the queue using the post-embedding algorithm described above and emitted events containing onset, bird ID, the low dimensional point, a letter representing the region it belonged to and the latency measured from the end of the segment until the event was emitted. The median latency over 3 million syllables was 1.089 s (percentiles: 5th=0.356, 25th=0.945, 75th=1.304, 95th=2.500). We found that 95 % of the segments were classified and the remaining 5 % were marked as unclassified.

Sequence events were generated, by event-processors in the cluster, based on the timing of the syllable events. If the onset of the next segment was within a window of 0.5 s after the offset of the previous, it was added to the sequence and otherwise it was assigned to a new sequence. Within 3 s after the end of a sequence an event was emitted containing onset, offset, bird id and the sequence.

To find the motif of the bird we formed a transition probability matrix based on the training data. Since the transitions in the motif were by far the most frequent, the syllables in the motif already stood out. Because the motif was repeated several times in a bout, they formed cycles in the transition matrix (Fig 4H). We found the cycle by starting from the globally most frequent transition and following the locally most frequent transitions until getting back to an already visited element. If the motif contained repeated syllables or if the bird sang a lot of introductory notes, there was a possibility for dead ends, but they could be detected and solved programmatically.

The birds often sing variations of the long motif so we found the ten most common substrings of the motif and looked for those in the sequence events. We counted the number of occurrences of each substring in the sequence and if a motif was present, we emitted a motif event (based on the most frequent substring in the sequence) containing onset, motif, number of occurrences and bird id. The motif detector was implemented as an event-processor running in the cluster. To verify the motif detector, an observer (LBL) looked at the spectrograms of all the motifs generated online during the two hours of experimentation for one bird (96 motifs) and confirmed that all of them were indeed motifs.

